# Yorkie and JNK revert syncytial muscles into myoblasts during Org-1 dependent lineage reprogramming

**DOI:** 10.1101/607820

**Authors:** Christoph Schaub, Marcel Rose, Manfred Frasch

## Abstract

Lineage reprogramming has become a prominent focus in research since it was demonstrated that lineage restricted transcription factors can be used *in vitro* for direct reprogramming [1]. Recently, we reported that the ventral longitudinal musculature (VLM) of the adult *Drosophila* heart arises *in vivo* by direct lineage reprogramming from alary muscles (AM), a process which starts with dedifferentiation and fragmentation of syncytial alary muscles into mononucleate myoblasts. Central upstream activators of the genetic program regulating the development of VLMs from alary muscles are the T-box factor Org-1 (*Drosophila* Tbx1) and the LIM homeodomain factor Tup (*Drosophila* Islet1) [2]. However, the events downstream of Org-1 and Tup that exert dedifferentiation and fragmentation of alary muscles have been unknown. In the present report, we shed light on the initiation of this first step of transdifferentiation and show that AM lineage specific activation of Yorkie (Yki), the transcriptional co-activator of the transcription factor Scalloped (Sd), has a key role in initiating AM lineage reprogramming. An additional necessary input comes from active dJNK signaling, which contributes to the inactivation of the Hippo kinase cascade upstream of Yki and furthermore activates dJun. The synergistic activities of the Yki/Sd and dJun/dFos (AP-1) transcriptional activator complexes in the absence of Hippo activity initiate AM dedifferentiation and lead to the expression of *Myc* and *piwi*, which are crucial for different aspects of AM transdifferentiation. Our results provide new insights into the mechanisms that mediate muscle lineage plasticity during a cellular reprogramming process occurring *in vivo*.

**Highlights:**

- Direct lineage reprogramming of alary muscles depends on Yorkie and JNK
- Yorkie and JNK mediate reversal of syncytial muscle cell fate
- Yki/Sd and AP-1 induce alary muscle dedifferentiation synergistically
- Yki dependent Myc induces and Piwi mediates reprogramming of alary muscles

## Results and Discussion

The formation of the VLMs of the adult *Drosophila* heart initiates with dedifferentiation and fragmentation of anterior larval syncytial alary muscles (AMs) into mononucleate myoblasts, called alary muscle derived cells (AMDCs). The AMDCs subsequently form syncytia and re-differentiate into the VLMs (Fig. 1A-C). In a recently performed alary muscle specific *in vivo* RNAi-screen (C.S. & M.F., unpublished data) we identified *scalloped* (*sd*) and *yorkie* (*yki*) as crucial genes for VLM formation. *org-1-GAL4* mediated knockdown of *sd* or *yki* within the alary muscles by either cell type specifically induced RNAi (Fig. S1B,C) or CRISPR (Fig. 1D,E), abolishes VLM formation and leads to the presence of *org-1-RFP* positive muscles that may represent remnants of the larval AMs. YAP, the mammalian homolog of Yki, can function as coactivator for several transcription factors including members of TEAD/TEF family. In *Drosophila,* Yki forms a complex with the single TEAD/TEF family member Sd that can activate transcriptional targets after nuclear translocation [3, 4]. Visualization of a GFP-tagged version of the endogenous protein clearly demonstrates that Sd is located in the nuclei of the syncytial *org-1-RFP* positive AM cells (Fig. 1F,F’). Although we were not able to detect nuclear GFP tagged Yki protein expressed under endogenous control (*yki::GFP*) in intact AMs, it does show a distinct nuclear accumulation during fragmentation of the AMs into *org-1-RFP* positive AMDCs (Fig. 1G,G’). Live imaging revealed that the phenotypes in *yki* knockdown backgrounds result from the inability of the AMs to dedifferentiate into AMDCs (movie S1). Thus, we propose that Yki as well as Sd are crucial for the initiation of AM transdifferentiation.

**Figure 1:**
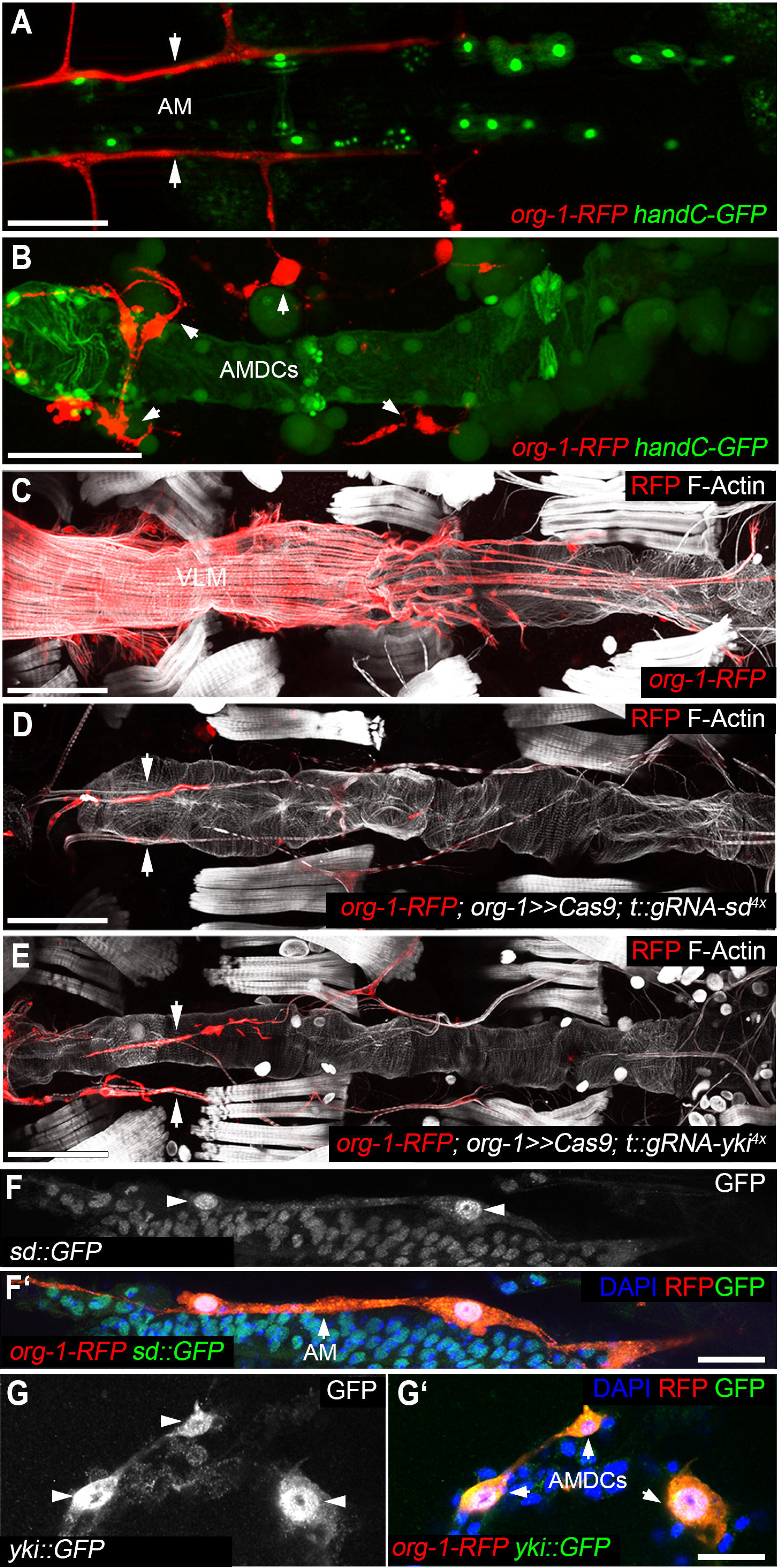
Scalloped (Sd) and Yorkie (Yki) are required for alary muscle dedifferentiation. **(A,B)** *Ex vivo* life imaging of a dissected white pupa carrying the *org-1-HN18-RFP* (*org-1-RFP*) and *hand-GFP* reporter constructs. **(A)** After the onset of metamorphosis *org-1-RFP* expression is initialized in the first three pairs of alary muscles (AM, arrows) that **(B)** initiate in stage P3 dedifferentiation and fragment into mononucleated progenitor like cells, the alary muscle derived cells (AMDCs, arrows). **(C)** *org-1-RFP* drives reporter expression in the ventral longitudinal musculature (VLM) attached to the adult heart. **(D,E)** Induction of CRISPR in the alary muscles with *HN39-org-1-GAL4* (*org-1-GAL4*) against **(D)** *sd* (*org-1*>>*Cas9; t::gRNA-sd*^*4x*^) or **(E)** *yki* (*org-1*>>*Cas9; t::gRNA-yki*^*4x*^) blocks VLM formation and prevents AM fragmentation (arrows). **(F,G)** Visualization of the *org-1-RFP* lineage marker and GFP tagged versions of *sd* (*sd::GFP*) and *yki* (*yki::GFP*). (F-F’) *sd::GFP* can be detected at the onset of metamorphosis in the nuclei (arrowheads) of the syncytial AM (arrow) whereas **(G-G’)** *yki::GFP* can be detected during induction of AM fragmentation in the nuclei (arrowheads) of the forming AMDCs (arrows). Scale bars in A-E: 100 μm. Scale bars in F-G: 10 μm. Actin is visualized with phalloidin. DNA visualized with DAPI.

The evolutionarily conserved Hippo pathway has been identified as one of the key inputs regulating Yki/YAP activity, particularly in the control of organ size via calibrating cell proliferation and apoptosis. Its core is represented by a kinase cascade consisting of Hippo (Hpo) [5–7] and Warts (Wts) that phosphorylates Yki, thereby inhibiting its nuclear translocation and its function as a transcriptional co-activator of Sd [3, 4, 8, 9]. To dissect a potential role of Hpo signaling during AM lineage reprogramming, we overexpressed various Hpo constructs (Fig. 2A) with *org-1-GAL4*. Strikingly, forced expression of an N-terminal fragment possessing the Ste20 kinase domain but not the regulatory domains (*UAS-Hpo*^*Δ*_*C*_^) completely abrogates AM transdifferentiation (Fig. 2B,D). Since this effect is not observed upon expression of the same deletion construct carrying an inactive Ste20 kinase domain (*UAS-Hpo*^*Δ*_*C.K71R*_^; Fig. 2E), of full-length Hpo (*UAS-Hpo*; Fig. 2F) or of its regulatory C-terminal part (*UAS-Hpo*^*Δ*_*N*_^; Fig. 2G) in the AMs, this strongly suggests that activated Hpo signaling blocks the induction of lineage reprogramming in the AMs. To prove that Hpo pathway mediated suppression of Yki function provokes the observed phenotypes in Hpo^Δ_C_^ overexpression backgrounds, we co-expressed it together with a phosphorylation resistant, constitutive active version of Yki (*UAS-yki*^*S168A*^). This leads to a significant rescue of VLM formation (p≤0.0001; Fig. 2B,H), thus strengthening our hypothesis that inactivation of the Hpo kinase cascade may lead to the activation of the Yki/Sd transcriptional activator complex that is critical for inducing AM reprogramming. Strikingly, by overexpressing Hpo-insensitive Yki^S168A^ in a genetic background in which we induced RNAi against *org-1* or its direct target *tup* [2], we observed that constitutively active Yki achieves a significant rescue of VLM formation (*org-1* p≤0.01, *tup* p≤0.0001; Fig. 2B,I-L). Taken together, these results are fully consistent with a mechanism in which Org-1 dependent inactivation of the Hippo pathway and concomitant activation of Yki/Sd target gene programs are needed to trigger alary muscle dedifferentiation into AMDCs.

**Figure 2:**
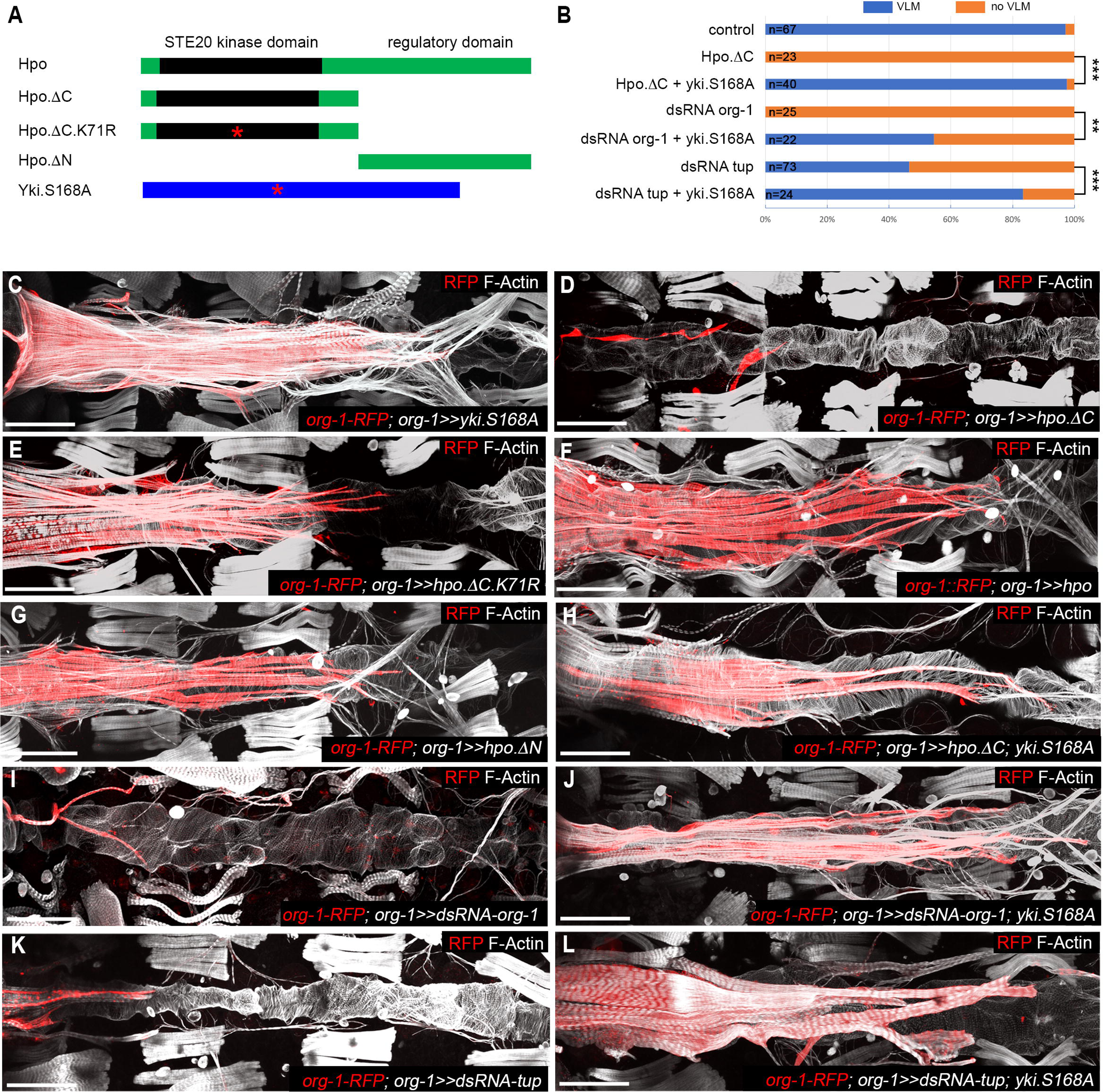
Org-1 dependent derepression of Yki is needed for alary muscle lineage reprogramming. **(A)** Schematic drawings of Hpo and Yki constructs used for overexpression. Asterisks indicate positions of amino acid exchanges. **(B)** Frequencies of observed VLM differentiation in the different genetic backgrounds. Co-expression of Yki^S168A^ can significantly rescue the phenotypes provoked by forced expression of Hpo^Δ_C_^ as well as the phenotypes induced by RNAi against Org-1 and Tup. **(C)** Forced overexpression of phosphorylation resistant Yki^S168A^ (*org-1*>>*yki.S168A*) with *org-1-GAL4* does not provoke a VLM phenotype. **(D)** Overexpression of the N-terminal part of Hpo containing a functional STE20 kinase domain (*org-1*>>*hpo.ΔC*) disrupts VLM formation and alary muscle fragmentation. **(E-G)** Forced overexpression with *org-1-GAL4* of **(E)** the N-terminal part of Hpo containing a dead STE20 kinase domain (*org-1*>>*hpo.ΔC.K71R*), *(F)* full length Hpo (*org-1*>>*hpo*) or **(G)** a N-terminally truncated version of Hpo (*org-1*>>*hpo.ΔN*) does not provoke a significant VLM phenotype. **(H)** VLM formation and alary muscle lineage reprogramming can be partially rescued by co-expression of Yki^S168A^ in Hpo^Δ_C_^ background (*org-1*>>*hpo.ΔC; yki.S168A*). **(I,K)** Induction of RNAi with *org-1-GAL4* against **(I)** Org-1 (*org-1*>>*dsRNA-org-1*) or **(K)** Tailup (Tup) (*org-1*>>*dsRNA-tup*) disrupts VLM formation and alary muscle lineage reprogramming. **(J,L)** Co-expression of Yki^S168A^ leads to the partial rescue of the phenotypes induced by the RNAi against either **(J)** Org-1 (*org-1*>>*dsRNA-org-1; yki.S168A*) or **(L)** Tup (*org-1*>>*dsRNA-tup; yki.S168A*). Actin is visualized with phalloidin. Scale bars: 100 μm.

In a first effort to explore possible links between Org-1 and Hippo signaling during lineage reprogramming, we tested additional known inputs into this pathway. In recent work it was proposed that aPKC signaling can regulate the nuclear outputs of the Hippo cascade either positively, namely by dislocating and inactivating the Hippo kinase [10], or in the case of mouse intestinal cells negatively, namely by directly phosphorylating YAP via the aPKC member PKCζ [11]. To test if aPKC activity plays a role in Hpo regulation during AM transdifferentiation we expressed a membrane tethered, dominant negative form of aPKC (*UAS-aPKC*^*CAAX-DN*^) as well as a constitutively active aPKC deletion construct (*UAS-aPKC*^*Δ*_*N*_^) via *org-1-GAL4*. Downregulation of aPKC activity does not have any impact on VLM formation, and thus does not appear to cause increased Hpo activity in this context. By contrast, the presence of the constitutively active form of aPKC abrogates VLM formation and AM lineage reprogramming, which could be due to increased Hpo and/or decreased Yki activity (Fig. 3B,C). In line with this possibility, the loss of VLM formation in the aPKC gain of function genetic background can be rescued significantly, albeit only partially, by the co-expression of phosphorylation resistant Yki (p≤0.001; Fig. 3D,G). These results support a model in which aPKC signaling is involved in the upstream regulation of Yki activity and needs to be inactive during AM transdifferentiation.

**Figure 3:**
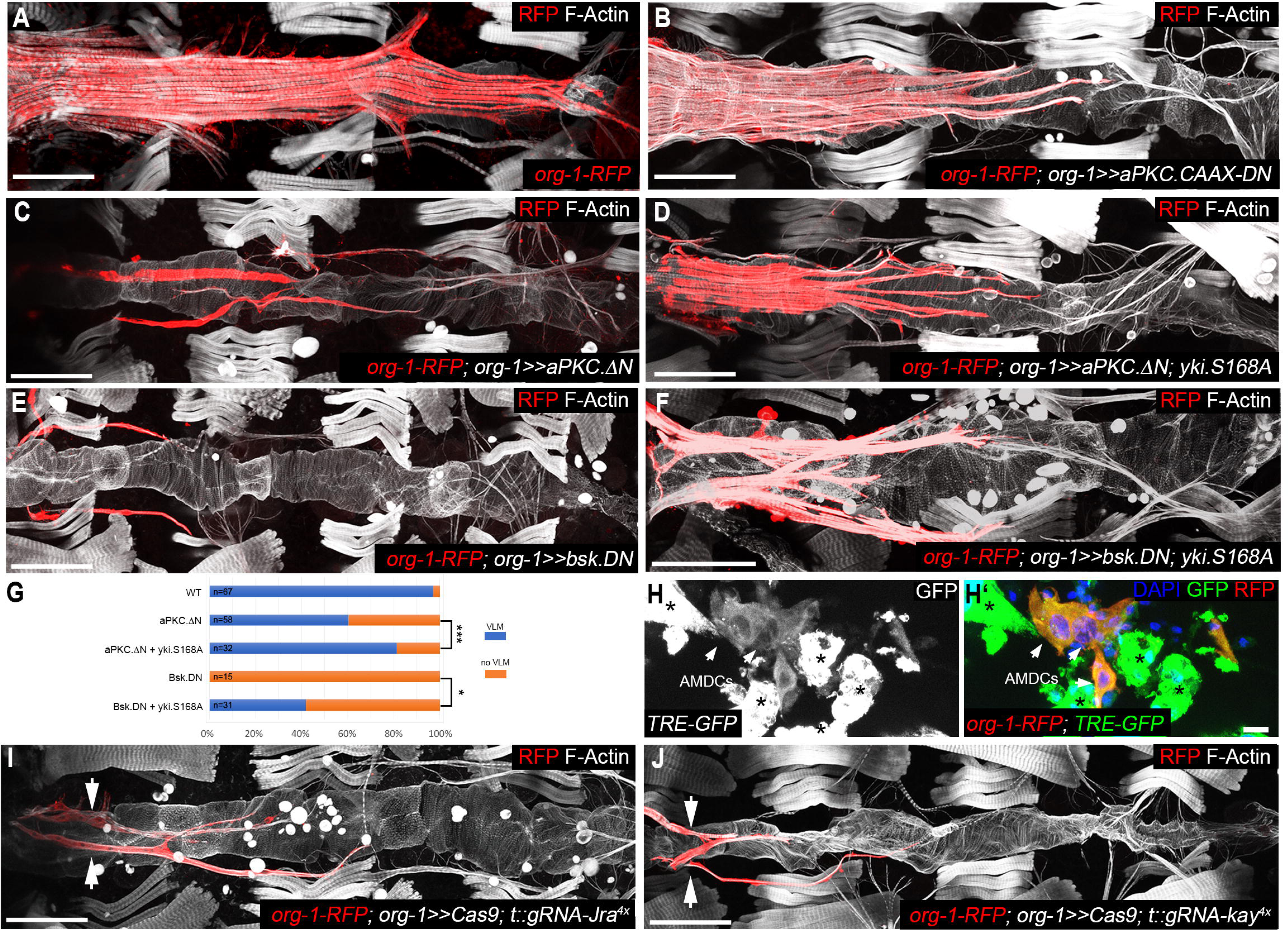
The transcriptional effectors of the Hippo and the dJNK pathways act synergistically to induce alary muscle fragmentation. **(A)** *org-1-RFP* drives reporter expression in the ventral longitudinal musculature. **(B)** Forced expression of a membrane tethered version of dominant negative aPKC (*org-1*>>*aPKC.CAAX.DN*) with *org-1-GAL4* has no effect on alary muscle reprogramming. **(C)** Induction of a constitutive active form of aPKC (*org-1*>>*aPKC.ΔN*) with *org-1-GAL4* disrupts VLM differentiation and AM transdifferentiation. **(D)** The “lost-VLM” phenotype in an aPKC gain of function background can be partially rescued by co-expression of phosphorylation resistant Yki^S168A^ (*org-1*>>*aPKC.ΔN; yki.S168A)*. **(E)** *org-1-GAL4* mediated induction of a dominant negative version of dJNK (*org-1*>>*bsk.DN*) leads to abolishment of VLM differentiation and alary muscle transdifferentiation. **(F)** The “lost-VLM” phenotype can only be weakly rescued by co-expression of Yki^S168A^ in dJNK^DN^ background (*org-1*>>*bsk.DN; yki.S168A*). **(G)** Frequencies of observed VLM differentiation in the different genetic backgrounds. Co-expression of Yki^S168A^ can significantly rescue the phenotypes provoked by forced expression of 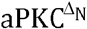 and dJNK^DN^ **(H,H’)** The *in vivo* AP-1 sensor *TRE-GFP* is activated in *org-1-RFP* positive AMDCs (arrows) and (even more strongly) in *org-1-RFP* negative apoptotic muscle cells (asterisks). **(I,J)** Induction of CRISPR in the alary muscles with *org-1-GAL4* against *(I) dJun* (*Jra)* (*org-1*>>*Cas9; t::gRNA-Jra*^*4x*^) or **(J)** *dFos* (*kay*) (*org-1*>>*Cas9; t::gRNA-kay*^*4x*^) blocks VLM differentiation and AM fragmentation (arrows). Scale bars in A-F&I-J: 100 μm. Scale bars in H: 10 μm. Actin is visualized with phalloidin. DNA is visualized with DAPI.

Since it has been shown that there is a functional link between the activation of JNK signaling and downregulation of Hpo pathway signaling [12], we next asked if Basket (Bsk, dJNK) activity is required for AM lineage reprogramming. To address this question, we overexpressed a dominant negative form of dJNK (*UAS-bsk*^*DN*^) in the alary muscles with *org-1-GAL4*, which resulted in the abolishment of VLM formation (Fig. 3E). Interestingly, co-expression of Bsk^DN^ and phosphorylation resistant Yki^S168A^ leads to a weak but significant rescue of VLM differentiation (p≤0.05; Fig. 3F,G), which indicates that active dJNK normally acts in promoting nuclear Yki, possibly also in this context via downregulating the Hippo pathway kinases. However, the incomplete rescue pointed towards additional inputs from dJNK signaling into AM lineage reprogramming. As a key component of the JNK pathway, dJNK phosphorylates the transcription factor Jun-related antigen (Jra, dJun) that subsequently together with Kayak (Kay, dFos) forms a heterodimeric protein complex, AP-1, which represents the transcriptional effector of JNK signaling [13]. By characterizing the expression pattern of an *in vivo* AP-1 sensor construct (*TRE-GFP*) during metamorphosis, we demonstrate *TRE-GFP* expression in *org-1-RFP* positive AMDCs (Fig. 3H,H’), thus implying that AP-1 targets are bound and activated during AM dedifferentiation. Moreover, downregulation of either *dJun* or *dFos* in the AMs by RNAi (Fig. S1D,E) or by inducible CRISPR (Fig. 3I,J) abolishes VLM formation and interferes with AM fragmentation, thus pointing to a crucial function of AP-1 during AM transdifferentiation.

Target genes of Yki have been mainly identified upon Yki hyperactivation or ectopic expression in imaginal discs. These experiments identified *Myc* as a transcriptional target of Yki, and it was shown that both work together in regulating proliferation and growth in epithelial imaginal cells [14, 15]. The analysis of the expression pattern of a GFP-tagged version of Myc under endogenous control (*Myc::GFP*) during AM lineage reprogramming revealed that, like *yki::GFP*, it can be detected in the nuclei of the AMDCs shortly after the initiation of syncytial AM fragmentation (Fig. 4B,B’). Downregulation of *Myc* via inducible RNAi or CRISPR during AM transdifferentiation strongly interferes with VLM formation and alary muscle reprogramming (Fig. S1F, Fig. 4D). Accordingly, live imaging demonstrated that upon RNAi mediated Myc knock-down alary muscle dedifferentiation and fragmentation is severely impaired (movie S2), pointing towards a crucial role for Myc in the initiation of AM reprogramming. Another target gene that has been identified upon Yki overexpression in *Drosophila* imaginal discs is *piwi* [16]. An endogenously controlled, GFP-tagged version of Piwi (*piwi::GFP*) is not yet detectable in syncytial AMs and AMDCs during AM fragmentation, but is expressed at high cytoplasmic levels during reprogramming of the AMDCs into the VLM progenitor cells (Fig. 4E-F’). Downregulation of *piwi* during AM transdifferentiation via inducible RNAi provokes loss of VLM formation (Fig. 4H). Live imaging revealed that upon *piwi* knock-down the anterior, *org-1-RFP* positive AMs do dedifferentiate and fragment into AMDCs but AM transdifferentiation is arrested at this stage (movie S3). This suggests an important role of Piwi during the reprogramming of the AMDCs into the progenitor cells of the VLMs. Notably, the induction of both *Myc::GFP* and *piwi::GFP* expression is suppressed by forced expression of constitutive active Hpo (*UAS-Hpo*^*Δ*_*C*_^) with *org-1-GAL4* in the AMs, thus confirming that they are essential Yki targets during AM lineage reprogramming (Fig. 4C-C’,G-G’).

**Figure 4:**
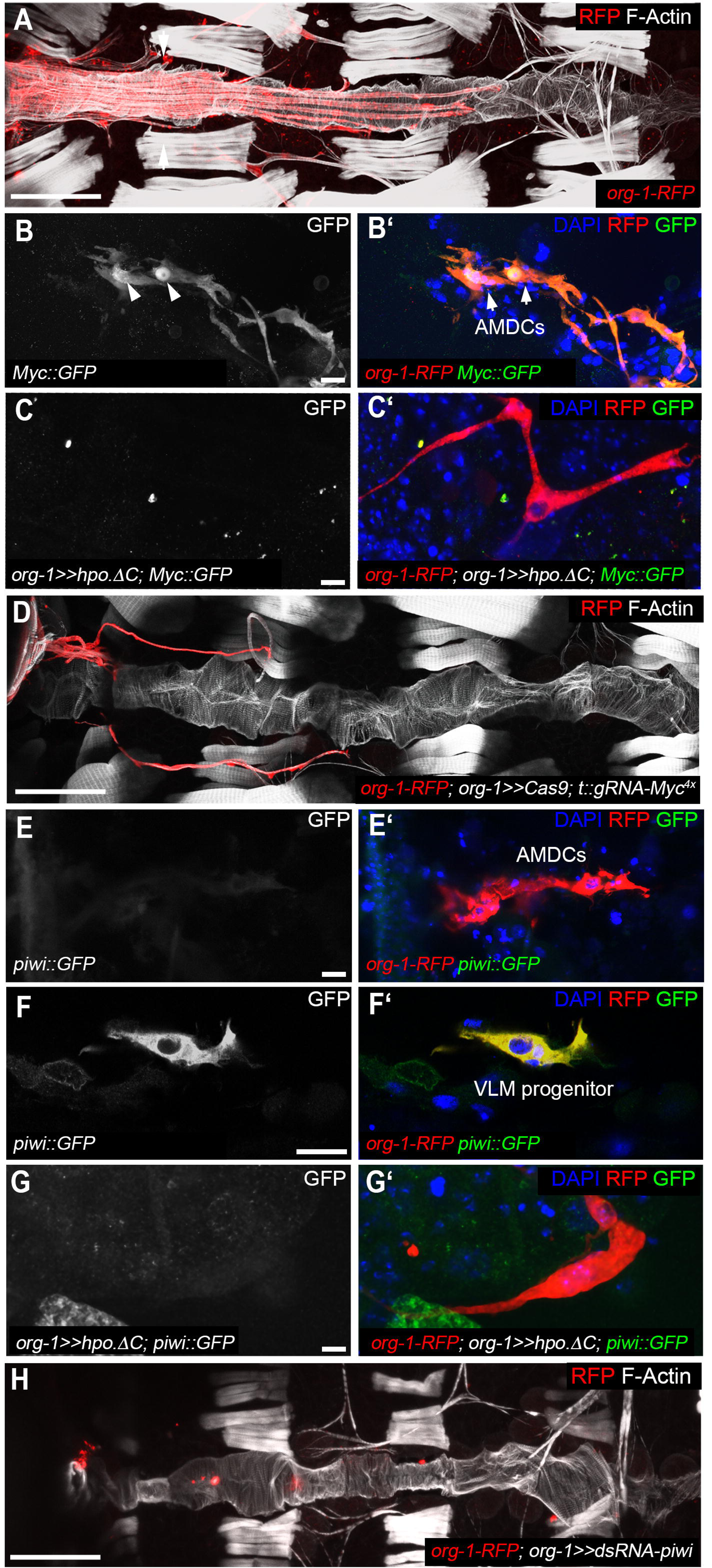
The Yki targets Myc and Piwi are indispensable for reprogramming of the alary muscle lineage. **(A)** *org-1-RFP* drives reporter expression in the ventral longitudinal musculature. **(B,C)** Expression of a GFP tagged *Myc* (*Myc::GFP*) **(B,B’)** can be detected in the nuclei (arrowheads) of the AMDCs (arrows) shortly after AM fragmentation, **(C,C’)** but is suppressed by forced overexpression of constitutive active Hpo with *org-1-GAL4* (*org-1*>>*hpo.ΔC*). **(D)** Induction of CRISPR in the alary muscles with *org-1-GAL4* against *Myc*(*org-1*>>*Cas9; t::gRNA-Myc*^*4x*^) abolishes VLM differentiation and blocks AM dedifferentiation. **(E,F)** Visualization of *org-1-RFP* and GFP tagged *piwi* (*piwi::GFP*) during AM lineage reprogramming. Whereas **(E,E’)** *piwi::GFP* cannot be detected in the forming AMDCs during fragmentation of the AMs, **(F,F’)** the VLM progenitor cells that arises from the AMDCs during reprogramming are clearly positive for cytoplasmic *piwi::GFP*. **(G,G’)** Forced overexpression of constitutive active Hpo with *org-1-GAL4* (*org-1*>>*hpo.ΔC*) abolishes *piwi::GFP* expression during AM transdifferentiation. **(H)** Induction of RNAi in the alary muscles with *org-1-GAL4* against *piwi* (*org-1*>>*dsRNA-piwi*) abolishes VLM differentiation. Scale bars in A, D, H,: 100 μm. Scale bars in B, C, E, F, G: 10 μm. Actin is visualized with phalloidin. DNA is visualized with DAPI.

Although syncytial muscles are often considered a paragon of terminally differentiated cells there are rare examples where skeletal muscle cells are induced to dedifferentiate and fragment into muscle progenitor cells, either by forced treatment or during a natural process as for the anterior *Drosophila* alary muscles [2, 17]. Our results herein reveal that effectors of the JNK and Hpo signaling pathways drive the lineage specific dedifferentiation and fragmentation of syncytial AM cells into mononucleate myoblasts. With regard to muscles, JNK function in *Drosophila* has been associated with myonuclear positioning and visceral muscle-intrinsic left-right asymmetry [18, 19] whereas Yki activity was described to be required for muscle growth during late pupal stages [20]. Both signaling pathways have also been implicated in dedifferentiation processes: active JNK in germ line cells [21] and Hippo (specifically, inactive Wts) in retinal cells [22]. We describe a novel, synergistic function of these two pathways in a specific type of muscle and connect them to lineage plasticity. Our data imply that the synergistic action of these two pathways in AM dedifferentiation is exerted predominantly in parallel at the level of their nuclear effectors, namely nuclear Yki/Sd, which requires inactivity of Hpo/Wts, and AP-1, which requires active JNK. Additionally, our data indicate that there is cross-regulation in which JNK also contributes to Yki activation, as has been described in studies on regeneration and size regulation in wing discs. As proposed for the wing disc, this input may involve inactivation of Wts upon JNK activation [12, 23]. The synergistic activities of the nuclear effectors of the Hippo and JNK cascades could operate directly at the level of joint target genes. Our genetic data and expression analysis implicate *Myc* and *piwi* as important examples of such targets, which in turn are required for mediating downstream effects of the Hippo and JNK cascades during this naturally occurring reprogramming process. In future studies, it should be examined whether these putative target genes contain combinatorial binding sites for Sd and AP-1.

Given that forced YAP expression can induce reprogramming of adult mammalian cells into cells with lineage progenitor characteristics [24], that the consensus motif for AP-1 and TEAD transcription factors are both significantly enriched in YAP-binding regions in cancer cell lines [25], and that the induction of pluripotent stem cells (iPS) requires the active JNK signaling pathway, c-Myc, and involves upregulation of *mili* (*piwil2*, a mouse homologue of *Drosophila piwi*) [26–28], we propose that the key components that drive AM muscle lineage reprogramming have broader, evolutionary conserved roles in mediating lineage plasticity. Specifically with regard to muscle tissues, it is worth noting in this context that in activated mouse satellite cells YAP expression surges, activated YAP protein promotes their proliferation and inhibits their differentiation, and conditional YAP knockout in these muscle stem cells curtails muscle regeneration [29, 30]. Likewise in the mouse heart, conditional depletion of the activity of the Hippo pathway kinases by various means, or expression of activated YAP, promotes heart regeneration beyond the early postnatal stages it is normally limited to. These effects involve dedifferentiation, reversal to a fetal gene expression program, and proliferation of cardiomyocytes [31]. Although the role of Hippo inactivation and Yki/YAP activation in tissue growth has been amply documented, these and other findings reinforce the notion that, depending on the context, this pathway can also exert cellular dedifferentiation. Of note, in the tissue context examined herein the dedifferentiated AMDCs do not proliferate; instead, upon being reprogrammed into founder cells of the VLMs they directly undergo myoblast fusion with fusion-competent myoblasts that have multiplied and migrated from lateral sources [2]. Nevertheless, the intriguing parallels in the roles of inactive Hippo/active Yki/YAP in promoting *Drosophila* alary muscle and mouse cardiomyocyte dedifferentiation, as well as in maintaining the dedifferentiated state of mouse muscle stem cells, warrant the search for additional similarities, including the potential involvement of regulatory inputs and outputs described in our current study in muscle and heart regeneration in the mouse and other vertebrate models.

Like Hippo inactivation, Myc function is mainly known for promoting proliferation and growth. However, besides these roles it has been connected to the induction of proapoptotic genes in a cell autonomous as well as a non-cell-autonomous manner [32]. Conversely, it can also exhibit anti-apoptotic and pro-autophagic effects in cancer cells [33]. We infer that Myc activation during AM lineage reprogramming is instrumental in initiating the mechanistic apparatus that executes the dedifferentiation and fragmentation process. This may include the induction of the nonapoptotic activity of caspases as well as of autophagic genes, which appear to be required for the disintegration of AM muscle components and syncytial fragmentation (C.S., unpublished data).

We show that cytoplasmic Piwi function is crucially required in the generated AMDCs to proceed with transdifferentiation. Cytoplasmic expression of PIWI proteins is associated with lineage restricted somatic stem cells from *Hydra* to mammals and there seems to be a strong correlation between PIWI expression and plasticity [34]. Accordingly, we envision that Piwi function in the AMDCs may be involved in the suppression of AM lineage specific programs. This could provide increased plasticity as a prerequisite for the reprogramming of AMDCs into VLM progenitor cells. Rather than being involved in the suppression of transposable elements it is more likely that Piwi functions through the degradation of cellular mRNAs during this process, which is also compatible with its observed cytoplasmic localization in the VLM progenitors. Precedence for this type of function exists for example in the Piwi mediated degradation of a large set of uniformly distributed maternal transcripts around the onset of zygotic transcription in the early *Drosophila* embryo [35, 36], the degradation of mRNAs (and lncRNAs) during consecutive steps of spermatogenesis in the mouse via different PIWI family members so that development can transition to the next steps of meiosis and spermiogenesis [36], and the degradation of cellular mRNAs via Piwi in somatic niche cells to advance their development during *Drosophila* oogenesis. Intriguingly, one important target of Piwi during this latter process is *dFos* mRNA, which needs to be destabilized for enabling the somatic niche cells to normally differentiate into follicle cells [37]. It is thus conceivable that in AMDCs, Piwi exerts a negative feedback and destabilizes *dFos* among other alary muscle or dedifferentiation-specific mRNAs to allow the transition towards the VLM progenitor fate.

Taken together, we suggest a mechanism in which Org-1/Tup driven lineage restricted activation of Yki as well as JNK in the alary muscles triggers a mechanism that executes cellular reprogramming and whose key components have evolutionary conserved roles. Further dissection of this transdifferentiation process occurring naturally *in vivo* will uncover additional components linking these regulatory pathways and mediating muscle cell plasticity, which may also shed more light on general mechanisms of lineage plasticity and cellular reprogramming.

## Acknowledgements

We are grateful to Johannes März for helping us with dissections. We thank the *Drosophila* community for supporting us with fly stocks, in particular Dirk Bohmann, Sonsoles Campuzano, Fillip Port and Jin Jiang as well as the Vienna *Drosophila* Resource Center and the Bloomington *Drosophila* Stock Center. This work was funded by grants of the Deutsche Forschungsgemeinschaft (DFG) to C.S (SCHA 2091/1-1) and M.F. (FR 696/6-1) and was supported by the Optical Imaging Centre Erlangen (O.I.C.E).

## Author contributions

Conceptualization, C.S. and M.F; Investigation, C.S. and M.R.; Writing – Original Draft, C.S.; Writing – Review & Editing, C.S and M.F.; Funding Acquisition, C.S. and M.F.; Supervision, C.S. and M.F.

## Materials & Methods

### CONTACT FOR REAGENT AND RESOURCE SHARING

Further information and requests for resources and reagents should be directed to and will be fulfilled by the corresponding authors, Manfred Frasch and Christoph Schaub.

### EXPERIMENTAL MODEL AND SUBJECT DETAILS

Individuals of *Drosophila melanogaster* were cultured using standard techniques at 25°C. Both female and male animals were used for the experiments.

### METHOD DETAILS

#### *Drosophila* Genetics

For the induction of UAS driven transgenes at 25 °C the following driver was used: *org-1-GAL4, org-1-RFP*. A complete list of the UAS lines used in this study can be found in the Key Resources Table.

To visualize protein activity during metamorphosis we obtained endogenously expressed, fluorescent protein forms of Sd [38], Yki [39], Myc [40] and Piwi [41] from the Bloomington *Drosophila* Stock Center as well as the Vienna Drosophila Resource Center. To create UAS inducible and t::gRNA array expressing transgenes we utilized the system described in [42]. We amplified PCR fragments with the respective primers (see Table S1 for information about the gRNAs and Table S2 for primer sequences used) and 1 ng pCFD6 (Addgene, Cat. No: 73915) as template utilizing Q5 Hot Start High-Fidelity 2X Master Mix (New England Biolabs, #M0494L). The PCR fragments were gel-purified using the QIAquick Gel Extraction Kit (Qiagen, Cat. No: 28704) and the respective constructs were assembled with BbsI-HF (New England Biolabs, Cat. No: 3539S) digested pCFD6 backbone using the NEBuilder® HiFi DNA Assembly Cloning Kit (New England Biolabs, Cat. No: R5520S), following the instructions provided by the manufacturers.

Transgenes of the respective pCFD6 constructs were generated by a commercial embryo injection service (BestGene Inc, Chino Hills, CA 91709, USA) at the (P{y[+t7.7]CaryP}attP2; 68A4, chr3L), (P{y[+t7.7]CaryP}attP40; 25C6, chr2L) and (PBac{y+-attP-9A}VK00027; 89E11, chr3R) landing sites utilizing the phiC31/attP/attB system [43].

#### Immunostaining and Imaging

Dissections of pupal stages until P4 (for pupal staging see [44]) were performed after prefixation with 3,7 % formaldehyde for 30 minutes with microsurgery scissors in toto. For pupae from P4 and later as well as for pharate adults the pupal cases were removed prior to prefixation and dissection. For antibody stainings the dissected animals were fixed in 3,7 % formaldehyde for 10 minutes, washed three times for 30 minutes in PBT, blocked in 10 % Bovine Serum Albumin (BSA) for 1 hour and incubated with the primary antibodies for two days at 4 °C. Primary antibodies, listed in the Key Resources Table, were diluted as follows: anti-GFP (1:1000) and anti-RFP (1:500). To visualize filamentous actin Phalloidin-Atto 647N (1:1000, Sigma-Aldrich, Cat. No: 65906) was added to the antibody solution. After the incubation with the primary antibodies the dissections were washed three times for 30 minutes in PBT and incubated with the respective secondary antibodies, listed in the Key Resources Table, in 1:200 dilution overnight at 4 °C. Finally, the stained animals were washed three times for 30 minutes in PBT and were embedded into Vectashield containing DAPI (Vectorlabs, Cat. No: H-1200). Confocal pictures were taken with a Leica SP5 II (20x/0.7 HC PL APO Glycerol, 63x/1.3 HC PL APO Glycerol). Projections were done with Leica Application Suite X (LAS X, Leica Microsystems).

#### *In vivo* time-lapse imaging

Pupae were aligned on a strip of double-faced adhesive tape connected to a slide, covered with a drop of Halocarbon oil 700 (Halocarbon Products Corp, Sigma Aldrich Cat. No: H8898) and a coverslip. Time-lapse series were acquired on a Leica SP5 II confocal system using a HC PL APO 10x/0.4 Air. Acquisition was done over a time course of 15-20 hours with the following settings: Pinhole 1.4 AU, 1.5x optical zoom, scan speed 200 Hz, line averaging 3, Z-stack about 20 sections with a step size of 3 μm and time intervals of 10 min per stack. Movies were generated using Leica Application Suite X (LAS X, Leica Microsystems).

### QUANTIFICATION AND STATISTICAL ANALYSIS

For the quantification of the VLM phenotype frequency living pharate stages of the respective genotype were aligned on a microscope slide. With the aid of a fluorescence equipped microscope (Nikon, Eclipse 80i) we scored phenotypes with fully differentiated VLM (proper aligned and elongated muscle fibres) versus phenotypes that showed abrogation of VLM differentiation (no aligned muscle fibres). Statistical analysis of the samples (Table S2) was performed with the Chi square test function of Microsoft Excel (Microsoft) and statistical significance was defined by p≤0.05.

### KEY RESOURCES TABLE

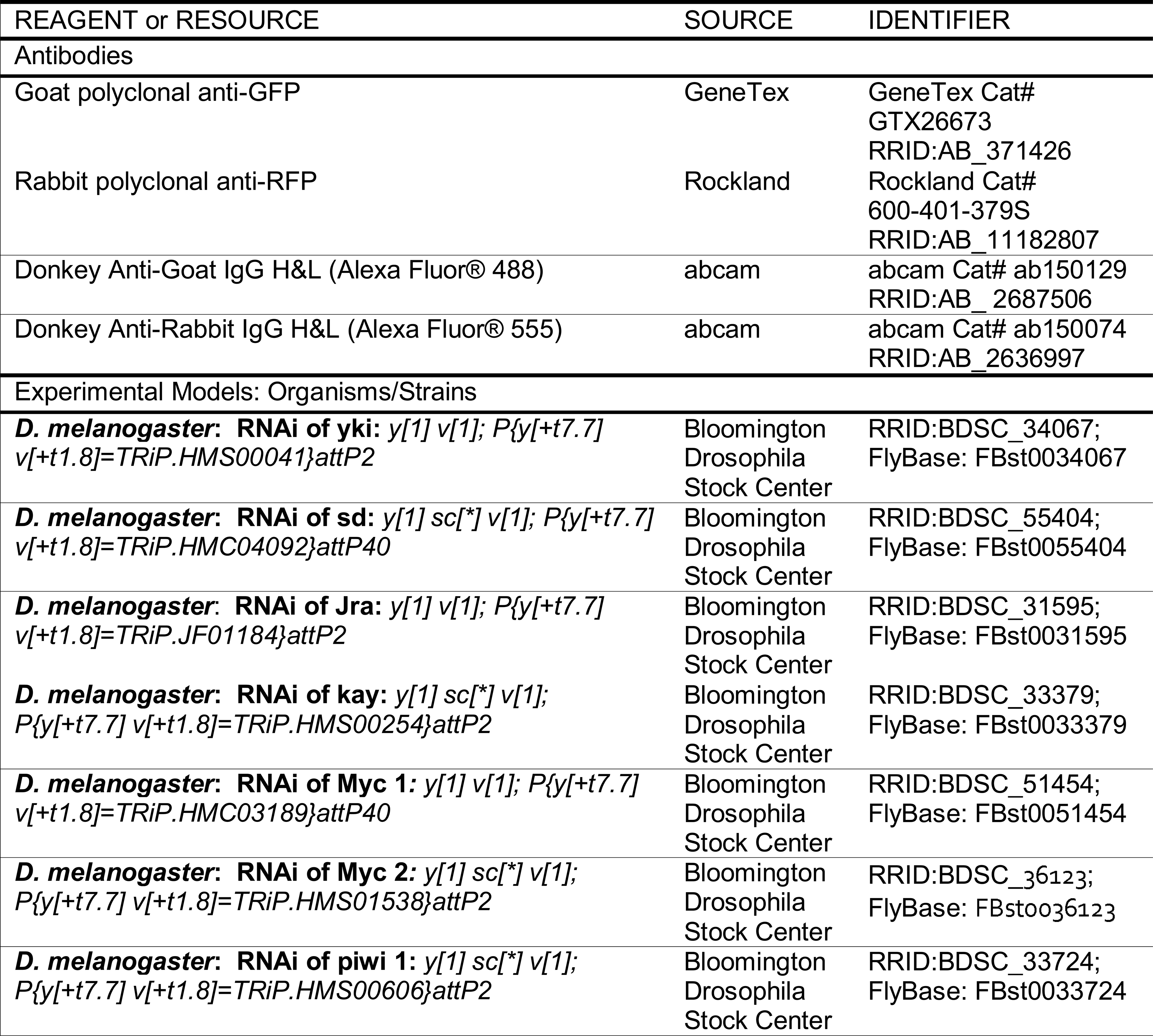
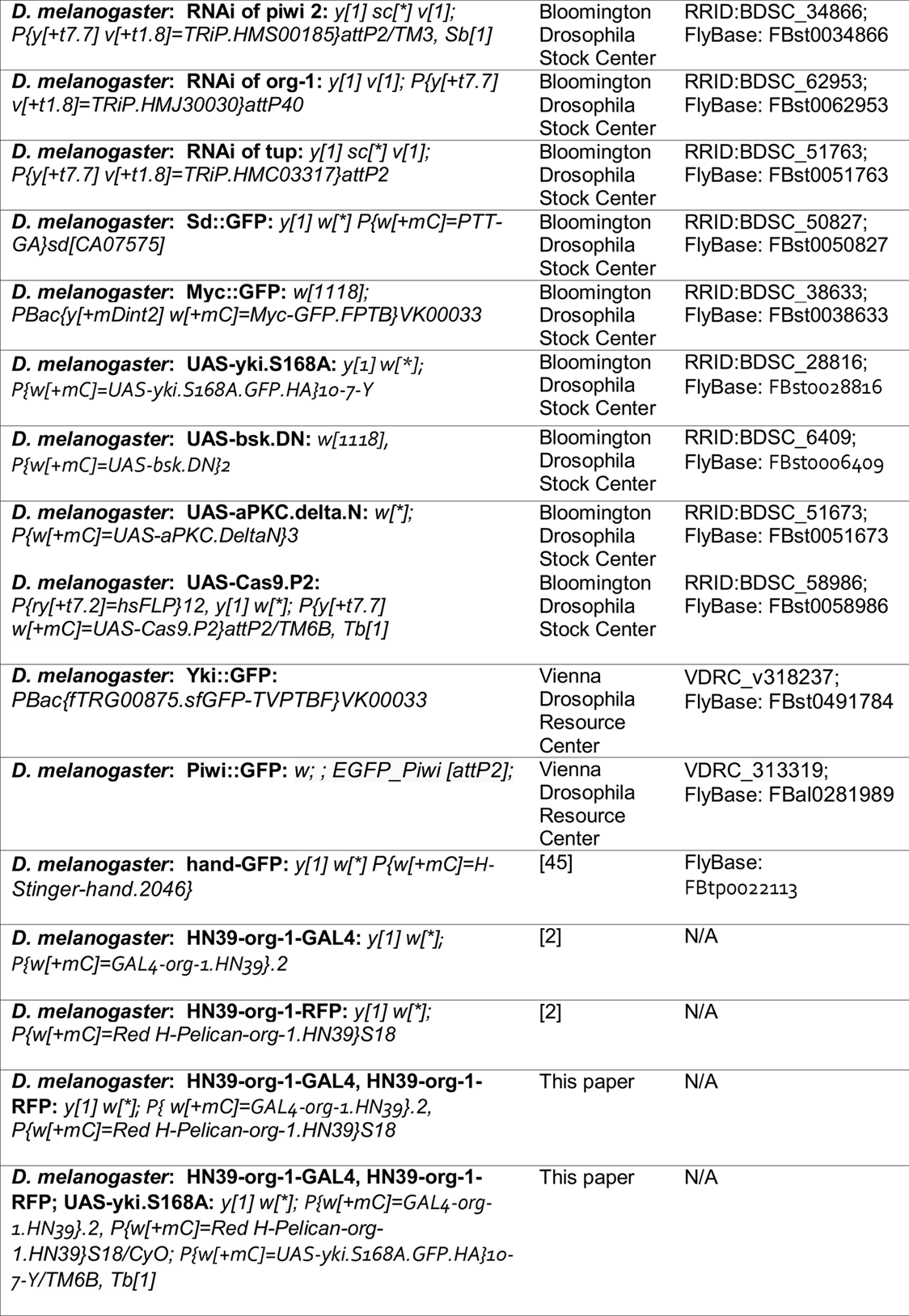
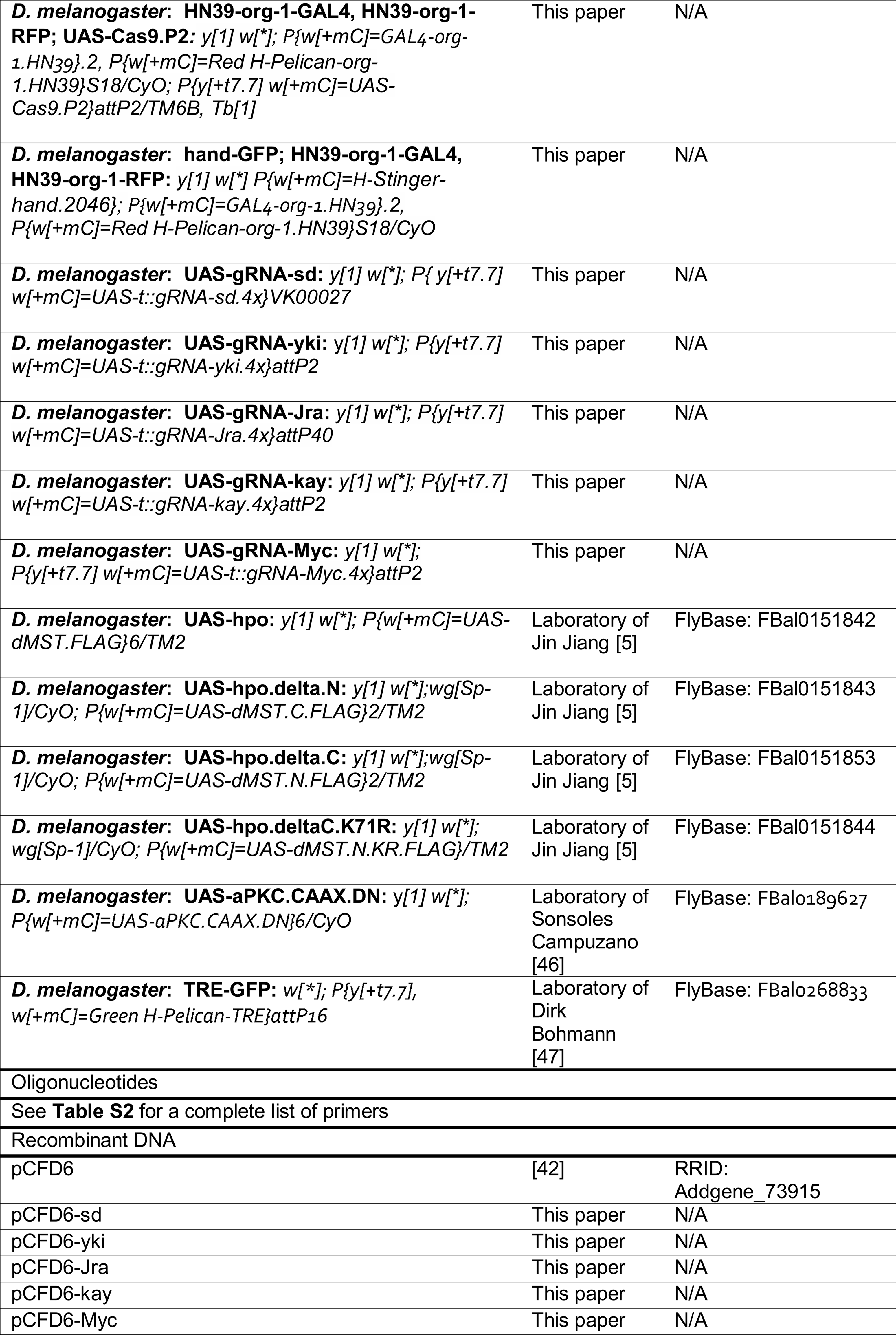

