## Supplemental Information for "Yorkie and JNK revert syncytial muscles into myoblasts during Org-1 dependent lineage reprogramming"

### Supplementary Figure

#### Figure S1. Induction of RNAi against the transcriptional effectors of Hpo and JNK as well as the Yki target Myc abolishes VLM differentiation

**(A)** *org-1-RFP*, as also shown in Fig. 1, drives reporter expression in the ventral longitudinal musculature (VLM) associated with the adult heart. **(B-F)** Induction of RNAi in the alary muscles with *org-1-GAL4* against **(B)** *sd* (*org-1>>dsRNA-sd*), **(C)** *yki* (*org-1>>dsRNA-yki*), **(D)** *Jra* (*org-1>>dsRNA-Jra*), **(E)** *kay* (*org-1>>dsRNA-kay*) and **(F)** *Myc* (*org-1>>dsRNA-Myc)* abolishes VLM formation and blocks AM dedifferentiation. Scale bars: 100 µm. Actin is visualized with phalloidin.

### Supplementary Movies

#### Movie S1.

Dorsal view of a pupa that carries *org-1-HN18-RFP* and *hand-GFP* and in which RNAi against *yki* was induced (*org-1-HN39-GAL4>>UAS-dsRNA-yki*). The movie starts at late stage P3 and shows that knock-down of *yki* prevents dedifferentiation of the alary muscles into mononucleate myoblasts, suggesting that Yki function is crucially required for the induction of this essential process during alary muscle transdifferentiation.

#### Movie S2.

Dorsal view of a pupa that carries *org-1-HN18-RFP* and *hand-GFP* and in which RNAi against *Myc* was induced (*org-1-HN39-GAL4>>UAS-dsRNA-Myc*). The movie starts at late stage P3 and shows that knock-down of *Myc* abolishes dedifferentiation of the alary muscles into mononucleate myoblasts, suggesting a pivotal role of the Yki target Myc in the genetic control of the alary muscle dedifferentiation process.

#### Movie S3.

Dorsal view of a pupa that carries *org-1-HN18-RFP* and *hand-GFP* and in which RNAi against *piwi* was induced (*org-1-HN39-GAL4>>UAS-dsRNA-piwi*). The movie starts at late stage P3 and shows that downregulation of *piwi* has no impact on the dedifferentiation of the alary muscles into mononucleate myoblasts but demonstrates that transdifferentiation is arrested under these conditions after alary muscle dedifferentiation. This suggests an essential role of the Yki target Piwi during the lineage reprogramming of the alary muscle derived myoblasts into the founders of the VLM during transdifferentiation.

### Supplementary Tables

| sd_gRNA_1 X:15820015..15820037 (+ strand) | CCGCTGATGCCGAAGGTGTATGG |
| --- | --- |
| sd_gRNA_2 X:15820071..15820093 (+ strand) | TTATCTATATATCCGCCGTGCGG |
| sd_gRNA_3 X:15820113..15820135 (+ strand) | TCCGACGAGGGTAAAATGTACGG |
| sd_gRNA_4 X:15819619..15819641 (+ strand) | CCGTGGACACCAGTGAATGCCGG |
| yki_gRNA_1 2R:24068081..24068103 (- strand) | ATTCGACAGCGTCCTGAATCCGG |
| yki_gRNA_2 2R:24068069..24068091 (+ strand) | GCTTGGCGTCACCCGGATTCAGG |
| yki_gRNA_3 2R:24067961..24067983 (- strand) | CGCCGACTCCACCTACGACGCGG |
| yki_gRNA_4 2R:24067953..24067975 (+ strand) | CTGGGAGCCCGCGTCGTAGGTGG |
| Jra_gRNA_1 2R:10097377..10097399 (- strand) | ACTGAAAGTCCAGTGACATGGGG |
| Jra_gRNA_2 2R:10097513..10097535 (- strand) | ACCGTCTTGGATGACAGATCCGG |
| Jra_gRNA_3 2R:10097421..10097443 (- strand) | GGGACGCTTGTTAGGATTCGGGG |
| Jra_gRNA_4 2R:10097345..10097367 (+ strand) | ATACCTAAAACCGAGCCCGTTGG |
| kay_gRNA_1 3R:29791413..29791435 (- strand) | TCGTCAGTGTGAGAACACTCTGG |
| kay_gRNA_2 3R:29791457..29791479 (- strand) | TGTGTCCTCGATGTTGCGCGTGG |
| kay_gRNA_3 3R:29791485..29791507 (- strand) | TCTGCGTGTCCGAGAGCAAGTGG |
| kay_gRNA_4 3R:29791510..29791532 (+ strand) | GATCGTGTGGCTGGTTGCGCGGG |
| Myc_gRNA_1 X:3375027..3375049 (- strand) | TGGTCGTCCATTATGGAATACGG |
| Myc_gRNA_2 X:3375124..3375146 (- strand) | CTTCTCGAGATCACTCTGAATGG |
| Myc_gRNA_3 X:3375537..3375559 (+ strand) | GAGGTCCATTTAATACCGCCCGG |
| Myc_gRNA_4 X:3375978..3376000 (- strand) | CTATCAGAGCCGGTCGTCGGCGG |

**Table S1.**

Sequences of gRNAs used for the knockdown of the respective genes. gRNAs have been obtained from <http://www.flyrnai.org/crispr/>

| sd_PCR1 fwd | CGGCCCGGGTTCGATTCCCGGCCGATGCACCGCTGATGCCGAAGGTGTAGTTTCAGAGCTATGCTGGAAAC |
| --- | --- |
| sd_PCR1 rev | CACGGCGGATATATAGATAATGCACCAGCCGGGAATCGAACC |
| sd_PCR2fwd | TTATCTATATATCCGCCGTGGTTTCAGAGCTATGCTGGAAAC |
| sd_PCR2 rev | TACATTTTACCCTCGTCGGATGCACCAGCCGGGAATCGAACC |
| sd_PCR3 fwd | TCCGACGAGGGTAAAATGTAGTTTCAGAGCTATGCTGGAAAC |
| sd_PCR3 rev | ATTTTAACTTGCTATTTCTAGCTCTAAAACGCATTCACTGGTGTCCACGGTGCACCAGCCGGGAATCGAACC |
| yki_PCR1 fwd | CGGCCCGGGTTCGATTCCCGGCCGATGCAATTCGACAGCGTCCTGAATCGTTTCAGAGCTATGCTGGAAAC |
| yki_PCR1 rev | GAATCCGGGTGACGCCAAGCTGCACCAGCCGGGAATCGAACC |
| yki_PCR2 fwd | GCTTGGCGTCACCCGGATTCGTTTCAGAGCTATGCTGGAAAC |
| yki_PCR2 rev | CGTCGTAGGTGGAGTCGGCGTGCACCAGCCGGGAATCGAACC |
| yki_PCR3f wd | CGCCGACTCCACCTACGACGGTTTCAGAGCTATGCTGGAAAC |
| yki_PCR3 rev | ATTTTAACTTGCTATTTCTAGCTCTAAAACCCTACGACGCGGGCTCCCAGTGCACCAGCCGGGAATCGAACC |
| Jra_PCR1 fwd | CGGCCCGGGTTCGATTCCCGGCCGATGCAACTGAAAGTCCAGTGACATGGTTTCAGAGCTATGCTGGAAAC |
| Jra_PCR1 rev | GATCTGTCATCCAAGACGGTTGCACCAGCCGGGAATCGAACC |
| Jra_PCR2 fwd | ACCGTCTTGGATGACAGATCGTTTCAGAGCTATGCTGGAAAC |
| Jra_PCR2 rev | CGAATCCTAACAAGCGTCCCTGCACCAGCCGGGAATCGAACC |
| Jra_PCR3 fwd | GGGACGCTTGTTAGGATTCGGTTTCAGAGCTATGCTGGAAAC |
| Jra_PCR3 rev | ATTTTAACTTGCTATTTCTAGCTCTAAAACACGGGCTCGGTTTTAGGTATTGCACCAGCCGGGAATCGAACC |
| kay_PCR1 fwd | CGGCCCGGGTTCGATTCCCGGCCGATGCATCGTCAGTGTGAGAACACTCGTTTCAGAGCTATGCTGGAAAC |
| kay_PCR1 rev | CGCGCAACATCGAGGACACATGCACCAGCCGGGAATCGAACC |
| kay_PCR2 fwd | TGTGTCCTCGATGTTGCGCGGTTTCAGAGCTATGCTGGAAAC |
| kay_PCR2 rev | CTTGCTCTCGGACACGCAGATGCACCAGCCGGGAATCGAACC |
| kay_PCR3 fwd | TCTGCGTGTCCGAGAGCAAGGTTTCAGAGCTATGCTGGAAAC |
| kay_PCR3 rev | ATTTTAACTTGCTATTTCTAGCTCTAAAACGCGCAACCAGCCACACGATCTGCACCAGCCGGGAATCGAACC |
| dMyc_PCR1 fwd | CGGCCCGGGTTCGATTCCCGGCCGATGCATGGTCGTCCATTATGGAATAGTTTCAGAGCTATGCTGGAAAC |
| dMyc_PCR1 rev | TTCAGAGTGATCTCGAGAAGTGCACCAGCCGGGAATCGAACC |
| dMyc_PCR2 fwd | CTTCTCGAGATCACTCTGAAGTTTCAGAGCTATGCTGGAAAC |
| dMyc_PCR2 rev | GGCGGTATTAAATGGACCTCTGCACCAGCCGGGAATCGAACC |
| dMyc_PCR3 fwd | GAGGTCCATTTAATACCGCCGTTTCAGAGCTATGCTGGAAAC |
| dMyc_PCR3 rev | ATTTTAACTTGCTATTTCTAGCTCTAAAACCCGACGACCGGCTCTGATAGTGCACCAGCCGGGAATCGAACC |

**Table S2.**

Sequences of the primers used for amplification of the fragments that has been utilized for the assembly of the respective CFD6 constructs (gRNA sequences indicated in red).

| **Genotype** | **VLM** | **no VLM** | **sum** | **p-value** |
| --- | --- | --- | --- | --- |
| ***control*** *org-1-GAL4, S18 x yw* | 67 | 2 | 69 |  |
| *org-1-GAL4, S18 x UAS-hpo. ΔC* | 0 | 23 | 23 |  |
| *org-1-GAL4, S18 x UAS-hpo. ΔC;  UAS-yki.S168A* | 39 | 1 | 40 | **0.0000000007** |
| *org-1-GAL4, S18 x UAS-dsRNA-org-1* | 0 | 25 | 25 |  |
| *org-1-GAL4, S18 x UAS-dsRNA-org-1;  UAS-yki.S168A* | 12 | 10 | 22 | **0.0105152459** |
| *org-1-GAL4, S18 x UAS-dsRNA-tup* | 34 | 39 | 73 |  |
| *org-1-GAL4, S18 x UAS-dsRNA-tup;  UAS-yki.S168A* | 20 | 4 | 24 | **0.0000000119** |
| *org-1-GAL4, S18 x UAS-aPKC. ΔN* | 35 | 23 | 58 |  |
| *org-1-GAL4, S18 x UAS-aPKC. ΔN;  UAS-yki.S168A* | 26 | 6 | 32 | **0.0000000002** |
| *org-1-GAL4, S18 x UAS-bsk.DN* | 0 | 15 | 15 |  |
| *org-1-GAL4, S18 x UAS-bsk.DN;  UAS-yki.S168A* | 13 | 18 | 32 | **0.0195502691** |

**Table S3.**

Frequencies of observed VLM phenotypes in the respective genetic backgrounds. P-values have been calculated utilizing the Chi square test.
