## Supplementary figures and images for "Yorkie and JNK revert syncytial muscles into myoblasts during Org-1 dependent lineage reprogramming"

### Figure S1

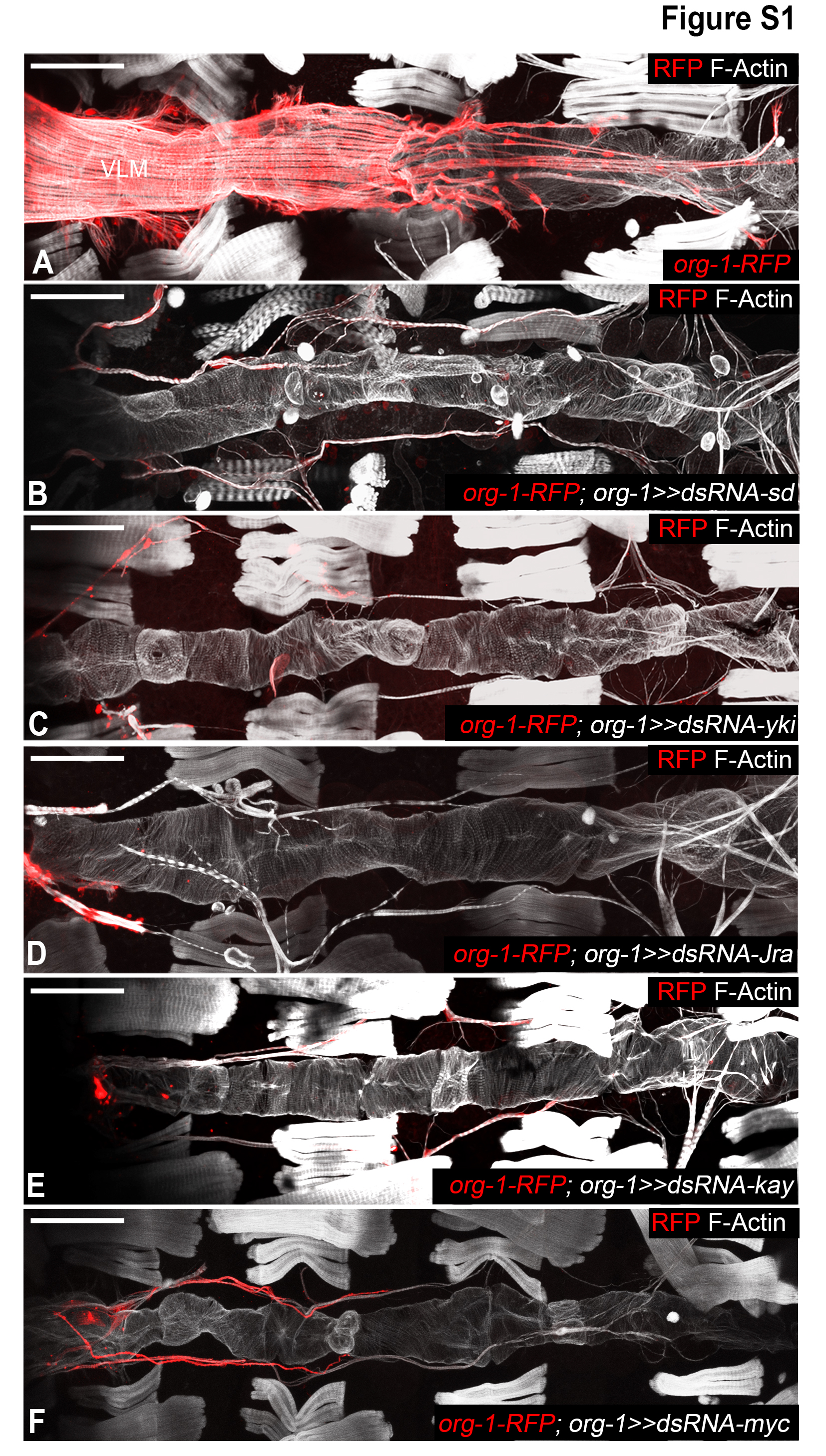
